# Inhibition of mTORC1 by rapamycin results in feedback activation of Akt^S473^ and aggravates hallmarks of osteoarthritis in female mice and non-human primates

**DOI:** 10.1101/2024.05.14.594256

**Authors:** Dennis M. Minton, Aditya R. Ailiani, Michael D.K. Focht, Mariana E. Kersh, Amy Li, Christian J. Elliehausen, Michelle M. Sonsalla, Dudley W. Lamming, Angela J. Marolf, Kelly S Santangelo, Adam B. Salmon, Adam R. Konopka

**Author notes:** These authors contributed equally (Dennis M. Minton, Aditya R. Ailiani). **Corresponding Author:** Adam R. Konopka, PhD, Division of Geriatrics and Gerontology Department of Medicine, University of Wisconsin-Madison, Geriatric Research, Education, and Clinical Center William S. Middleton Memorial Veterans Hospital.

## Abstract

**Purpose:** Genetic deletion of mTOR has protected against post-traumatic osteoarthritis (OA) in male mice, however, effects of pharmacological mTOR-inhibition are equivocal and have not been tested in aging models nor in female subjects. Therefore, the goal of this study was to determine if mTOR-inhibition by rapamycin can modify OA pathology in aging non-human primates and female mice.

**Methods:** Common marmosets were administered oral rapamycin (1mg/kg/day) or vehicle starting near mid-life until death. Five-month-old, female C57BL/6J mice were treated with vehicle or rapamycin (IP, 2mg/kg, 3x/week) for 8-weeks following non-invasive ACL rupture. Knee OA pathology was assessed via microCT and histology. Phosphorylation of mTORC1 (p-RPS6^S235/36^) and mTORC2 (p-Akt^S473^, p-NDRG1^T638^, p-PKCα^T348^) substrates were evaluated via western blot in articular cartilage, meniscus, and/or infrapatellar fat pad. ATDC5 cells were cultured with rapamycin to determine time and dose effects on mTORC1/2 signaling.

**Results:** In marmosets, rapamycin did not impact age-related radiographic OA severity or cartilage pathology but increased medial meniscus calcification and lowered lateral tibia subchondral thickness, particularly in females. In female mice, rapamycin worsened ACLR-induced meniscus calcification and cartilage pathology. In marmoset and mouse joint tissues, rapamycin inhibited mTORC1 and increased p-Akt^S473^ but not p-NDRG1^T638^ or p-PKCα^T348^. This mTOR signaling pattern was replicated in ATDC5 cells during exposure to low concentrations of rapamycin.

**Conclusions:** Rapamycin attenuated mTORC1 signaling with feedback activation of Akt^S473^ in articular cartilage, meniscus, and/or infrapatellar fat pad and was accompanied by deleterious effects on meniscus calcification and/or cartilage pathology in female mice and common marmosets.

## INTRODUCTION

Increasing age and previous joint injury are the greatest risk factors for osteoarthritis (OA). OA further increases the age-related risk of multi-morbidity including cardiovascular disease, type 2 diabetes, dementia, frailty, and mobility limitations (1). With no available disease modifying therapies for OA, the economic and societal impacts of OA are expected to rapidly inflate with population aging. Critical barriers to the development of disease modifying therapies for human OA are 1) an incomplete mechanistic understanding of chondrocyte biology and OA progression and 2) lack of models that faithfully recapitulate human OA.

The mechanistic target of rapamycin (mTOR) is a nutrient- and growth factor-sensitive protein kinase which exists as two distinct protein complexes, mTORC1 and mTORC2 (2). mTORC1 acts on processes including protein translation through its effector RPS6, and mTORC2 acts on Akt, the SGK-NDRG1 axis, and PKC family members to relay insulin/growth factor signals, control cytoskeletal remodeling and promote cell survival (3). Phosphorylated mTORC1/2 substrates are abundant in OA cartilage from humans, dogs, and rodents (4–7). Increasing signaling through mTORC1 (6), and the mTORC2 substrate Akt (5), can induce and/or exacerbate OA in mice. Genetic deletion of the entire mTOR kinase has protected against surgically-induced OA in young male mice (7), however, pharmacological mTOR inhibition strategies have shown more equivocal results. Several studies report protective effects of mTOR-inhibitors in male models of experimental OA (4,7–9), while we and others have found no effect or worsened OA after treatment with the mTOR-inhibitor rapamycin (10–12). This lack of consensus warrants further investigation.

In preclinical animal models, rapamycin extends lifespan in both sexes and delays several age-associated pathologies (13–16). To determine the translational potential of these beneficial effects, rapamycin is now being tested in common marmosets, the smallest anthropoid non-human primate (17–19). Recently, we identified that the common marmoset replicates many aspects of human joint tissue organization, develops hallmark OA pathologies in an age-related fashion, and show a greater propensity of females to develop OA (20). Therefore, the marmoset represents an advantageous model to bridge the gap between rodents and humans.

Here we present findings to indicate rapamycin may worsen medial meniscus calcification and lead to subchondral bone loss, particularly in female marmosets. While marmosets provide high translational potential they come with significant heterogeneity. To corroborate these findings in more well controlled models, we tested the influence of mTOR inhibition by rapamycin on OA pathology in female mice after ACL rupture and determined dose- and duration-dependent effects of rapamycin on mTOR signaling *in vitro*.

## MATERIALS AND METHODS

### Common marmoset study overview

Common marmosets (n=65) were housed and maintained using previously published husbandry guidelines at the Barshop Institute for Longevity and Aging Research at UT Health San Antonio (19). Twenty-four (14M/10F) marmosets received eudragit-encapsulated rapamycin (1mg/kg/day) dissolved in yogurt. The control group consisted of 28 marmosets (14M/14F) that received yogurt vehicle with empty eudragit as previously published (19) and an additional 13 marmosets (7M/6F) from the general population that did not receive eudragit vehicle. All husbandry was identical, and there were no differences in OA between the vehicle control and age-matched general population marmosets. Marmosets in the control group for this study were previously published to characterize OA pathology in the common marmoset (20). Upon natural death or compassion euthanasia, hind limbs were either fixed in formalin (n=64) or frozen (n=19) and shipped to University of Wisconsin-Madison.

### Evaluation of OA in common marmosets

Marmoset samples were processed and evaluated via microCT and histopathology as described in detail in our previous publication (20). For convenience, these details are also provided in **Supplementary Methods**.

### Mouse use and drug administration

All mouse experiments were approved by the Institutional Animal Care and Use Committee (IACUC) at University of Wisconsin-Madison and/or the William S Middleton Memorial Veterans Hospital. All mice were acclimated to housing for at least 1-week before procedures. Female C57BL/6J mice (n=24) were purchased from Jackson Laboratory (Bar Harbor, ME) and were housed at Wisconsin Institute for Medical Research and the Waisman Center at the University of Wisconsin-Madison. Mice were group housed under 12-hour light/dark cycle and were provided ad libitum access to food and water. Rapamycin (LC Labs) was reconstituted 40mg/mL in sterile 100% ethanol then diluted in vehicle (5% Tween-80, 5% PEG-40, in 0.9% NaCl) for injection. Mice received intra-peritoneal (IP) injection of rapamycin (2mg/kg) or vehicle with equivalent ethanol concentration 3x/week (M/W/F) for 8-weeks.

For a separate experiment, male mice were purchased from Jackson Laboratory at 8-weeks of age. Everolimus (LC Labs) was aliquoted into individual vials containing 4.05mg per vial then each vial was reconstituted for every other injection by first dissolving the everolimus in 80μl of 100% ethanol before dilution in vehicle (same as above) for injection. Mice received IP injections of everolimus (3mg/kg) every other day for 5 weeks.

### Non-invasive ACL rupture

At 5-months of age, female mice underwent a unilateral ACL rupture (ACLR). The morning of the procedure, mice were injected with 0.6mg/kg extended-release buprenorphine. Mice were anesthetized via isoflurane until unresponsive and maintained for the duration of the procedure. Left hind limbs were subjected to ACLR as previously described (21) and right limbs were used as internal contralateral controls. Briefly, left limbs were positioned in a MARK10 ESM303 mechanical testing unit (Copiague, NY, USA) and subjected to 0.6N pre-load. To ensure complete stretch-relaxation of the ACL, limbs were manually adjusted until secure in knee/heel restraints and pre-load force was stable. Pre-load was then maintained at 0.6N for 30-seconds. A compressive load was applied to achieve a displacement rate of 1mm/s across a total displacement range of 1.67mm, and ACL rupture was evident by auditory cues and/or a drop on force-displacement curve. Mice were returned to normal housing once fully recovered from anesthesia as evident by grooming and normal ambulatory behavior. Mice were monitored for 3-days following the procedure.

### Sacrifice and tissue collection

For sacrifice of rapamycin-treated mice, mice were injected with rapamycin or vehicle, fasted overnight (12-14-hours), and re-fed ∼1.5-hours prior to sacrifice via cervical dislocation, as this approach has previously been shown to reveal rapamycin-induced mTORC2-inhibition (22). Hind limbs were collected into 10% neutral buffered formalin for 48-hours and transferred to 70% ethanol. Cartilage was collected from the proximal humerus, snap frozen in liquid nitrogen, and stored at −80C.

For everolimus-treated mice, mice were injected with everolimus or vehicle ∼20-hours prior, fasted overnight, and sacrificed via cervical dislocation. Cartilage was collected from the proximal humerus, snap frozen in liquid nitrogen, and stored at −80C.

### microCT acquisition and analysis

Mouse contralateral and ACLR hind limbs were scanned submerged in 70% ethanol using an MILabs U-SPECT/CT, achieving an isometric voxel resolution of 10-microns. In Amira (Thermo), scans were binarized at identical thresholds across samples, joint tissues (menisci, fabellae, patella) were segmented, and calcified volumes were quantified using the Material Statistics function.

### Histopathology

For evaluation of OA pathology in mice, a Modified OARSI scoring system which has been previously used for ACLR-induced PTOA (23) was employed (**Table S1**). During evaluation, “tibial degeneration” was considered erosion past the tidemark on the posterior aspect of the tibia, and “staining present on the femur” was considered to be the presence of toluidine-blue-stained non-calcified cartilage on the femoral condyle. In some cases, scores were assigned in increments of 0.5.

### Fasting blood glucose

Fasting glucose was measured prior to mice receiving their third weekly dose of rapamycin. Following a 4-6-hour fast initiated at ∼7am, blood glucose was measured from the tail vein using a Contour glucometer and glucose strips (Bayer).

### Cell culture

ATDC5 cells were purchased from Sigma Aldrich and maintained in DMEM/F12 50:50 mix with L-glutamine (Gibco), 5% fetal bovine serum, and 1% penicillin/streptomycin at 37C and 5% CO_2_. Cells were seeded in 6-well plates, and Rapamycin or Torin-1 (LC Labs) were reconstituted in DMSO and added to maintenance media at the indicated concentrations and durations, achieving a DMSO concentration of 0.25% v/v which was applied to vehicle controls. For time-course experiments, doses were staggered to ensure the time from the last media change to lysis was consistent between groups. All experiments were performed with at least n=3 per condition.

### Western blotting

Tissue samples were homogenized in TRIzol and protein was extracted as previously described (24). Cell cultures were lysed in radioimmunoprecipitation buffer (RIPA) with protease and phosphatase inhibitors on for 10-minutes on ice, centrifuged at 10,000xG for 10-min at 4C, and supernatant was collected.

Protein quantification was performed via Pierce BCA Assay (Thermo), and samples were prepared in reducing conditions with B-mercaptoethanol in 4x Laemmli buffer (BioRad). Equal amounts of protein were then separated on 7% or 4-15% gradient TGX pre-cast gels (BioRad) and transferred to Nitrocellulose membranes. Membranes were stained with Ponceau S and imaged using a BioRad ChemiDoc or Cytivia ImageQuant 800. Membranes were then blocked in 5% bovine serum albumin in TBST followed by incubation with the following primary antibodies diluted in blocking buffer: p-S235/36-RPS6 (#4858), RPS6 (#2217), p-S473-Akt (#4060), Akt (#4961), p-T346 NDRG1 (#3217), and p-T638-PKCα (#9375) all from Cell Signaling and respective HRP-conjugated anti-rabbit (#7074, Cell Signaling) or anti-mouse (ab6728, Abcam) secondary antibodies. Membranes were imaged after a 5-minute incubation in SuperSignal Pico or Femto chemiluminescent substrates (Fisher), or a 50:50 mix of the two, depending on target abundance. After imaging, membranes were stripped in Restore Stripping Buffer (Fisher) for 15-minutes and re-probed for additional targets. Densitometric analysis was performed in Image Lab (BioRad). p-RPS6^S235/36^ and p-Akt^S473^ were expressed relative to their respective total protein while p-NDRG1^T346^ and p-PKCα^T638^ were expressed relative to Ponceau S.

### Statistical analysis

For analysis of common marmoset data, two primary approaches were used to assess the impact of age, sex, and rapamycin treatment on OA outcomes. First, we assessed the impact of age as a continuous variable using linear regressions for control and rapamycin-treated marmosets when pooled and stratified by sex. Trendlines were compared to determine if rapamycin impacted the age-related trajectory of dependent variables. Second, we performed pooled comparisons of control and rapamycin-treated geriatric (>8-years-old) marmosets using unpaired t-tests or Mann-Whitney tests, depending on normality of data. These comparisons were also made when pooling and stratifying by sex but did not include adults (<8-years-old) due to small number of rapamycin-treated adults. One control marmoset was deemed a statistical outlier by Grubbs test (alpha=0.01) and was not included in analysis of meniscus volume.

For mouse studies, longitudinal data and data including contralateral and ACLR limbs were analyzed using 2-way repeated measures ANOVA with the Holm-Sidak multiple comparison test. For pooled comparisons between groups, unpaired t-tests, Mann-Whitney tests, or one-way ANOVA were used, depending on number of groups and normality of data.

## RESULTS

### Rapamycin treatment duration and physical characteristics

Average treatment durations for marmosets pooled and stratified by age-group and sex are shown in **Table S2**. On average, marmosets were treated with rapamycin for 2.1±1.5 years. Peak body mass and blood glucose were also measured on a subset of marmosets as shown in **Table S3** and were not significantly different between control and rapamycin.

### Rapamycin did not modify radiographic or histological cartilage pathology

Representative microCT 3D reconstructions of knee joints closest to group median are shown in **Figure 1A**. Total OA scores were positively related to age when pooled and stratified by sex, with the exception of rapamycin-treated female marmosets (**Figure 1B**). However, no differences were found between trendlines of control and rapamycin-treated. Similarly, in geriatric marmosets, microCT OA scores were not different between control and rapamycin-treated marmosets when pooled and stratified by sex (**Figure 1C**). No differences between treatment groups were found for individual scoring criteria (data not shown).

**Figure 1:**
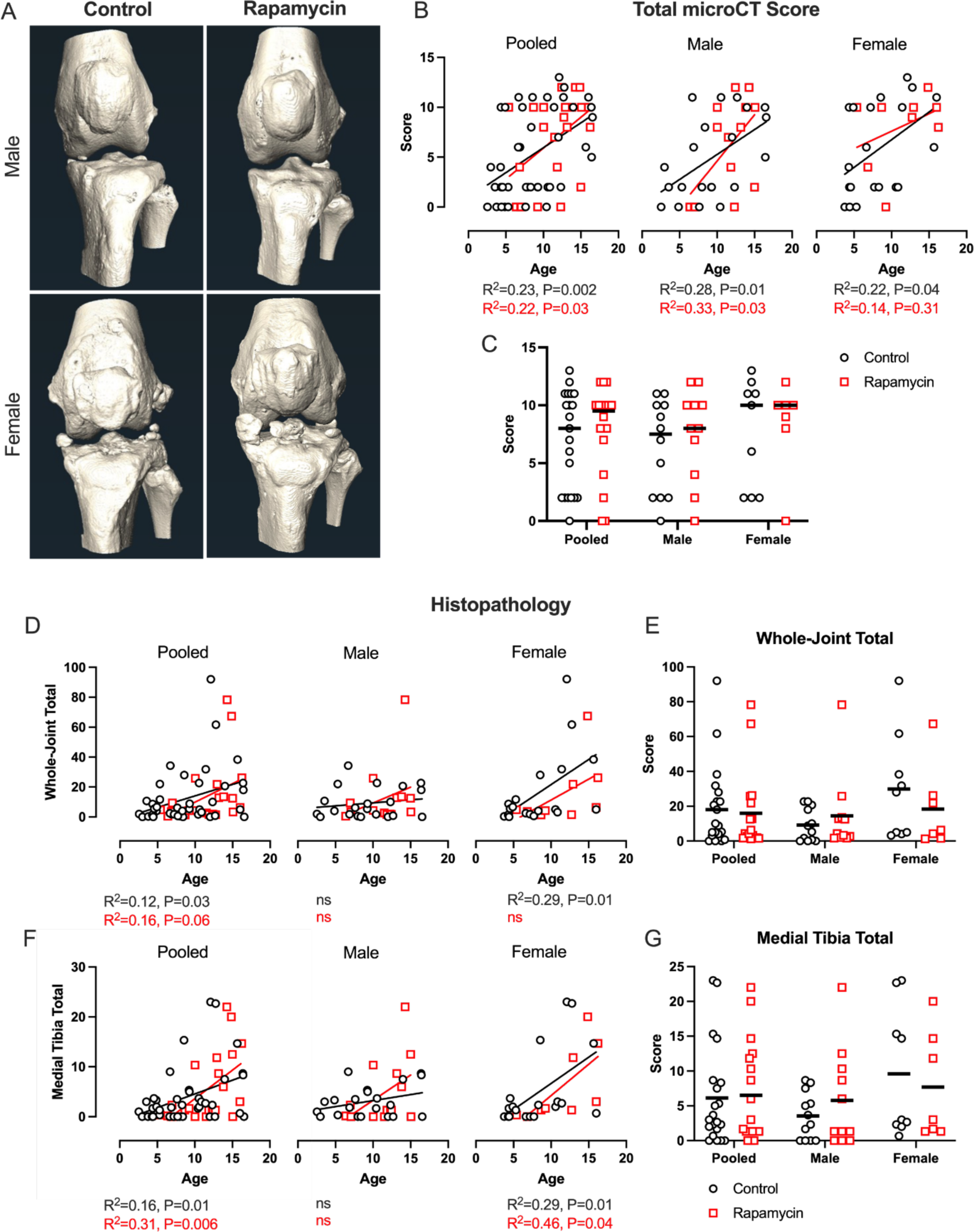
Rapamycin does not attenuate age-related radiographic OA severity in common marmosets. **A**) Representative 3D reconstructions of male and female control and rapamycin-treated marmoset knee joints. **B**) Linear regression of total microCT score against age are shown for control and rapamycin-treated marmosets when pooled and stratified by sex. Linear regression results are shown beneath figures; no trendline differences were found between groups. **C**) Total microCT scores of geriatric marmosets were compared between treatment groups when pooled and stratified by sex. **D**) Linear regressions against age of summed whole-joint Modified Mankin scores from control and rapamycin-treated marmosets are shown when pooled and stratified by sex. No trendline differences were found between treatment groups. **E**) Whole-joint scores of geriatric marmosets were compared between treatment groups when pooled and stratified by sex. **F**) Linear regressions against age of total Modified Mankin scores from the medial tibia of control and rapamycin-treated marmosets are shown when pooled and stratified by sex. No trendline differences were found between treatment groups. **G**) Medial tibia scores of geriatric marmosets were compared between treatment groups when pooled and stratified by sex. Data were analyzed via Pearson’s R (B,D,F) or Mann-Whitney tests (C,E,G) and are presented as individual data points with trendlines, median (C), or mean (E,G).

Cartilage pathology assessed by modified Mankin scoring is shown in **Figure 1 D-F**. While cartilage pathology in the whole joint and medial tibial plateau were positively associated with age, trendlines were not different between control and rapamycin-treated marmosets when pooled or stratified by sex. There were no differences between control and rapamycin-treated geriatric marmosets for whole joint or medial tibial plateau. No treatment effects were found in other joint compartments or for individual scoring criteria (data not shown).

### Greater meniscus calcification in female rapamycin-treated marmosets

Representative images of median medial meniscus calcification volumes are shown in **Figure 2A**. Linear regressions (**Figure 2B**) demonstrate an age-related increase in meniscus calcification in control (R^2^=0.28, P<0.001) and rapamycin-treated (R^2^=0.25, P=0.02) marmosets, with a steeper trendline slope in the rapamycin group. When stratifying by sex, trendlines were nonsignificant and superimposable between groups for males. By contrast, meniscus calcification was highly related to age in control (R^2^=0.66, P<0.001) and rapamycin-treated females (R^2^=0.63, P=0.01), with a higher slope in the rapamycin group (P=0.002). In geriatric marmosets, medial meniscus calcification was not significantly different between groups when pooled or stratified by sex, despite rapamycin-treated female marmosets having three-fold greater median meniscus calcification volume than controls (**Figure 3C**). Together, these data indicate rapamycin may worsen the age-related increase in meniscus calcification in marmosets, particularly in females.

**Figure 2:**
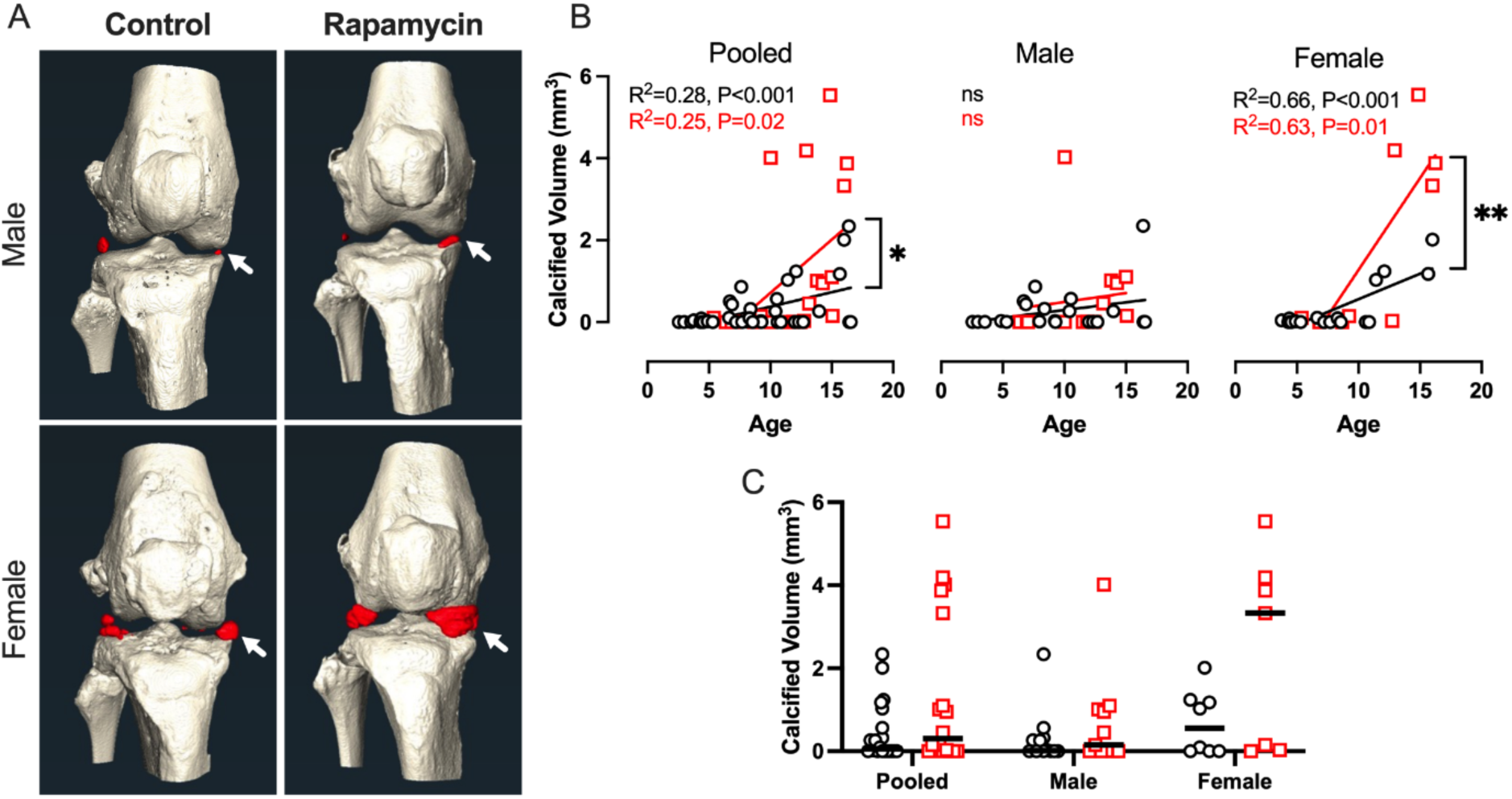
Calcification of the medial meniscus in control and rapamycin-treated marmosets. **A**) 3D microCT reconstructions representative of median values of medial meniscus calcification for geriatric male and female control and rapamycin-treated marmosets. Meniscus calcification is highlighted in red, and the medial meniscus is denoted with a white arrow. **B**) Linear regressions between age and medial meniscus calcification volume are shown for pooled and stratified sexes. Linear regression results and differences between trendlines are shown within each graph, when significant. **C**) Pairwise comparisons were performed between geriatric control and rapamycin-treated marmosets when pooled and stratified by sex. Are presented as trendlines or median with individual data points and were analyzed by linear regression (B) or Mann-Whitney tests (C).

**Figure 3:**
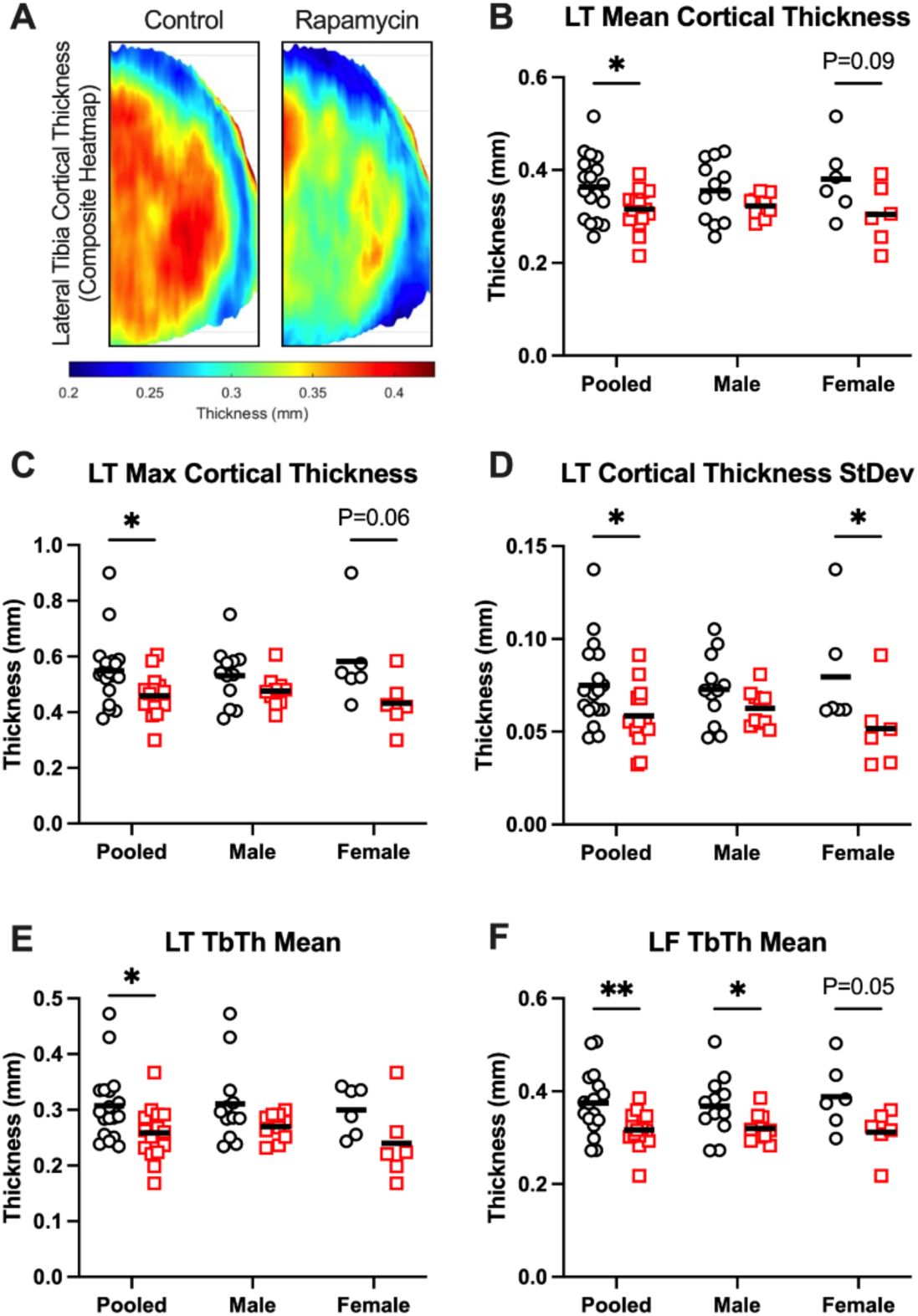
Subchondral bone architecture is altered by rapamycin treatment. **A**) Composite cortical thickness heatmaps from control and Rapamycin-treated marmosets, scale bar beneath. **B)** Mean, **C**) max, and **D**) standard deviation of lateral tibia cortical bone thickness were all lower in Rapamycin-treated marmosets than controls. **E**) Trabecular thickness in the lateral tibia and **F**) lateral femur were also lower in Rapamycin-treated marmosets than controls. L=lateral, T=tibia, F=femur. Data are presented both pooled and stratified by sex. Treatment effects were assessed by unpaired t-tests or Mann-Whitney tests, depending on normality of data. Data are shown as mean with individual data points. *P<0.05, **P<0.01

### Rapamycin decreased marmoset lateral tibial subchondral bone thickness

Composite heatmaps of cortical bone thickness from the lateral tibia of control and rapamycin-treated marmosets are shown in **Figure 3A**. Lateral tibia cortical thickness (mean, max, and standard deviation) were all lower (P<0.05) in geriatric rapamycin-treated marmosets versus control when sexes were pooled (**Figure 3B-D**). These effects were driven by females, and no significant differences were found for males. Trabecular thickness was also lower in rapamycin-treated geriatric marmosets in the lateral tibia (**Figure 3E**, P<0.05) and lateral femur (**Figure 3F**, P<0.01) when sexes were pooled. In the lateral femur, decreased trabecular thickness was seen in both males (P<0.05) and females (P=0.05). No treatment effects were seen in other joint compartments, and no significant differences between trendline slopes were seen between control and rapamycin-treated marmosets (data not shown).

### ACL rupture parameters and physical characteristics of female mice

Due to the heterogeneity of aging primates, we sought to investigate if the deleterious effects of rapamycin on hallmarks of OA observed in female marmosets were apparent in a more controlled OA model. Because female mice have traditionally shown relatively low levels of age-related OA, we employed a non-invasive ACL-rupture model to induce post-traumatic OA followed by treatment with rapamycin (2mg/kg, 3x/week) or vehicle.

During the ACLR procedure, the ACL failed at an average load of 11.65N (95%CI 10.87, 12.44) and 1.175mm (95%CI 1.064, 1.286) of displacement. Mice were randomized to ensure equivalent ACLR conditions between groups (**Figure S1A-B**). Bodyweight increased during the study (**Figure S1C**, time effect P<0.0001) and was not significantly different between groups at any timepoint measured. Consistent with notion that rapamycin disrupt glucose homeostasis (25), fasting blood glucose was higher in rapamycin-treated mice at 2-, 4-, and 8-weeks of treatment versus vehicle control (**Figure S1D**, treatment effect P<0.0001).

### ACLR-induced meniscus calcification and cartilage pathology were exacerbated in rapamycin-treated mice

Representative images of toluidine blue-stained sections from contralateral and ACLR limbs of vehicle and rapamycin-treated mice are shown in **Figure 4A**. Modified OARSI scoring revealed main effects for ACLR (P<0.0001) and an ACLR x treatment interaction (**Figure 4B**, P=0.003). Further, modified OARSI scores from the ACLR limb were significantly higher in rapamycin-treated mice than vehicle (P=0.0005).

**Figure 4:**
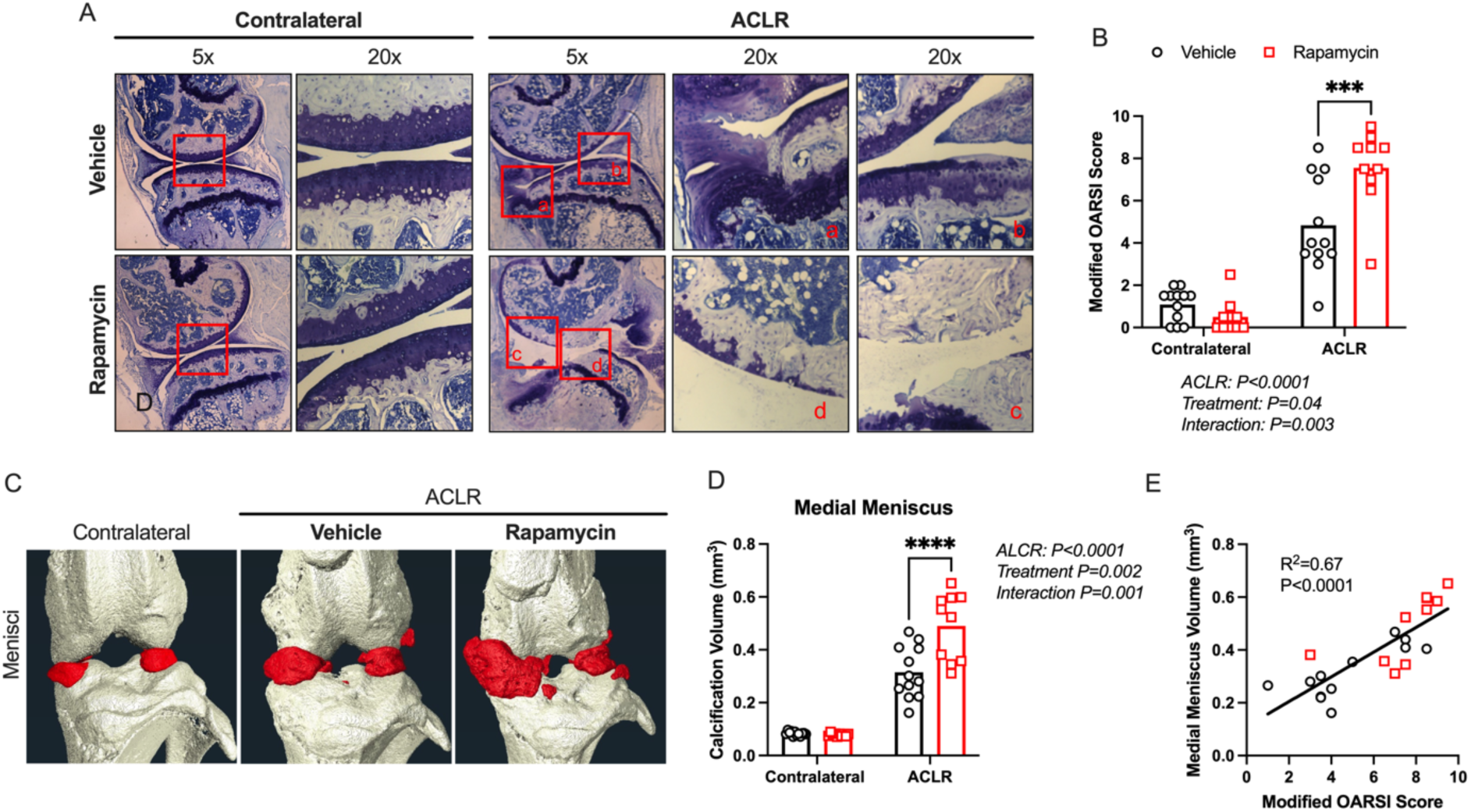
Increased ACLR-induced cartilage degeneration and meniscus calcification in rapamycin-treated mice. **A**) Representative toluidine-blue stained histology images from the medial compartment of contralateral and ACLR limbs of vehicle and rapamycin-treated mice. **B**) Images were quantified using the Modified OARSI scoring system tailored to the ACLR OA model. **C)** 3D reconstructions of microCT scans representative of a median contralateral limb and ACLR limbs from vehicle and rapamycin-treated mice. Regions of calcified meniscus are highlighted in red. **D**) Quantified meniscus calcification volumes from contralateral and ACLR knees. **E**) Linear regression between Modified OARSI scores and medial meniscus calcification volumes. Data are presented as mean with individual data points and were analyzed using 2-way repeated measures ANOVA with Sidak’s multiple comparison test, with ANOVA effects reported on each figure panel. N=10-12 per group, ***P<0.001, ****P<0.0001.

Representative images displaying meniscus calcification from a healthy contralateral limb, vehicle-treated ACLR, and rapamycin-treated ACLR limbs are shown in **Figure 4C**. In the medial meniscus, rapamycin exacerbated ACLR-induced calcification (**Figure 4D**, ACLR effect P<0.0001, ACLR x treatment P=0.001). Modified OARSI scores were also strongly correlated with medial meniscus calcification volume (**Figure 4E**; R^2^=0.67, P<0.0001).

### Rapamycin inhibits mTORC1 and increases Akt^S473^ phosphorylation in marmoset and mouse joint tissues

Akt signaling is elevated in human OA cartilage, and overactivation of Akt induces cartilage degeneration in mice (5). In non-articular cell types, low doses or acute exposure to rapamycin can activate Akt through alleviation of mTORC1-dependent negative feedback on insulin/IGF receptor signaling (26,27). It has been speculated that this may limit therapeutic efficacy or even contribute to more severe OA during rapamycin treatment (28). Consistent with this notion, oral rapamycin treatment inhibited the relative phosphorylation of the mTORC1 effector, p-RPS6^S235/236^, and increased the phosphorylation of the mTORC2 effector, Akt^S473^ in the tibio-femoral articular cartilage, meniscus, and infra-patellar fat pad in common marmosets (**Figure 5A, B**).

**Figure 5:**
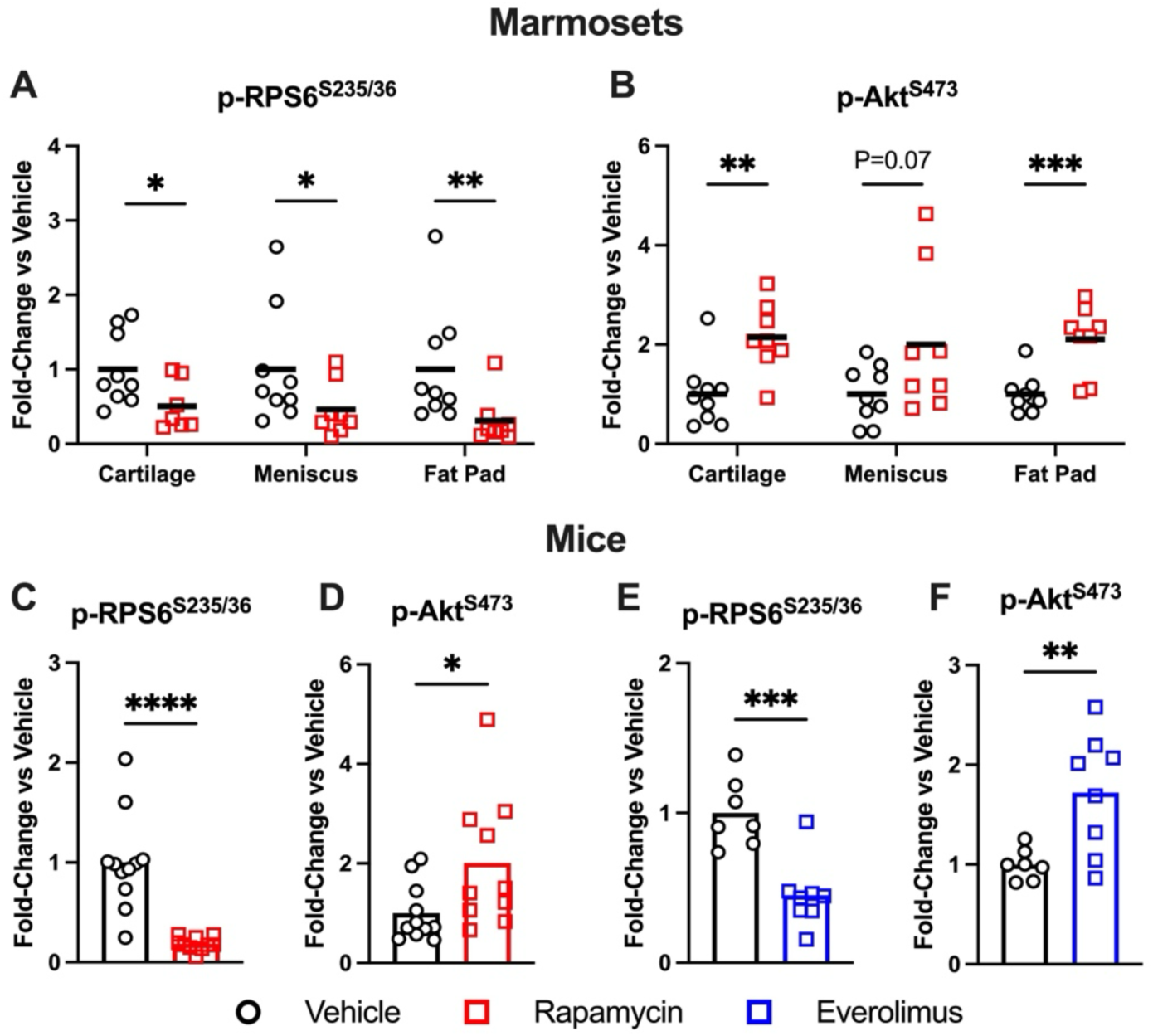
Treatment with mTOR-inhibitors increases signal through p-Akt^S473^ in marmoset and mouse joint tissues. **A)** Densitometric analysis of p-RPS6^S235/236^ normalized to total RPS6 and **B**) p-Akt^S473^ normalized to total Akt from control and rapamycin-treated marmoset articular cartilage, meniscus, infrapatellar fat pad. **C**) Densitometric analysis of p-RPS6^S235/36^ normalized to total RPS6 and **D**) p-Akt^S473^ normalized to total Akt from glenohumeral cartilage of female mice treated with rapamycin (2mg/kg, 3x/week) or vehicle for 8-weeks. **E**) Densitometric analysis of p-RPS6^S235/236^ normalized to total RPS6 and **F**) p-Akt^S473^ normalized to total Akt from glenohumeral cartilage of male mice treated with everolimus (3mg/kg every other day) or vehicle for 5-weeks. N=7-9/group (A-B), 10-12/group (C-D), or 7-8/group (E-F). Data are shown as mean with individual data points and were analyzed by student’s t-test. *P<0.05, **P<0.01, ***P<0.001, ****P<0.0001.

Next, we evaluated mTOR signaling in cartilage from the glenohumeral joint of female mice subjected to ACLR with or without rapamycin treatment. Relative phosphorylation of RPS6^S235/236^ was significantly decreased by rapamycin, indicating mTORC1 inhibition (**Figure 5C**, P<0.0001). Interestingly, this was partially driven by increased total protein abundance in the rapamycin-treated group. Conversely, relative Akt^S473^ phosphorylation was increased by rapamycin (**Figure 5D**, P<0.05).

To determine whether these effects are also seen in male mice during systemic treatment with an mTOR-inhibitor, we performed western blotting on glenohumeral cartilage from male mice receiving the rapamycin analog, everolimus (3mg/kg every other day, IP), or vehicle for 5-weeks. Similarly, we found everolimus decreased p-RPS6^S235/236^ **Figure 5E**, P<0.001) and increased p-Akt^S473^ (**Figure 5F**, P<0.01) versus vehicle. Together, these data demonstrate small molecules across the mTOR-inhibitor drug class result in mTORC1 inhibition with increased p-Akt^S473^ in joint tissues from common marmosets and mice.

### Alternative mTORC2 substrates are inhibited or unaffected by rapamycin in marmoset and mouse joint tissues

In addition to Akt, mTORC2 can also impact phosphorylation of NDRG1 and PKCα in other cell and tissue types. Therefore, we evaluated p-NDRG1^T346^ and p-PKCα^T638^ to determine if other mTORC2 substrates were similarly upregulated by rapamycin in joint tissues. p-NDRG1^T346^ was not significantly affected in marmoset joint tissues (**Figure 6A**), however, p-PKCα^T638^ was significantly decreased by rapamycin in marmoset cartilage and fat pad (**Figure 6B**).

**Figure 6:**
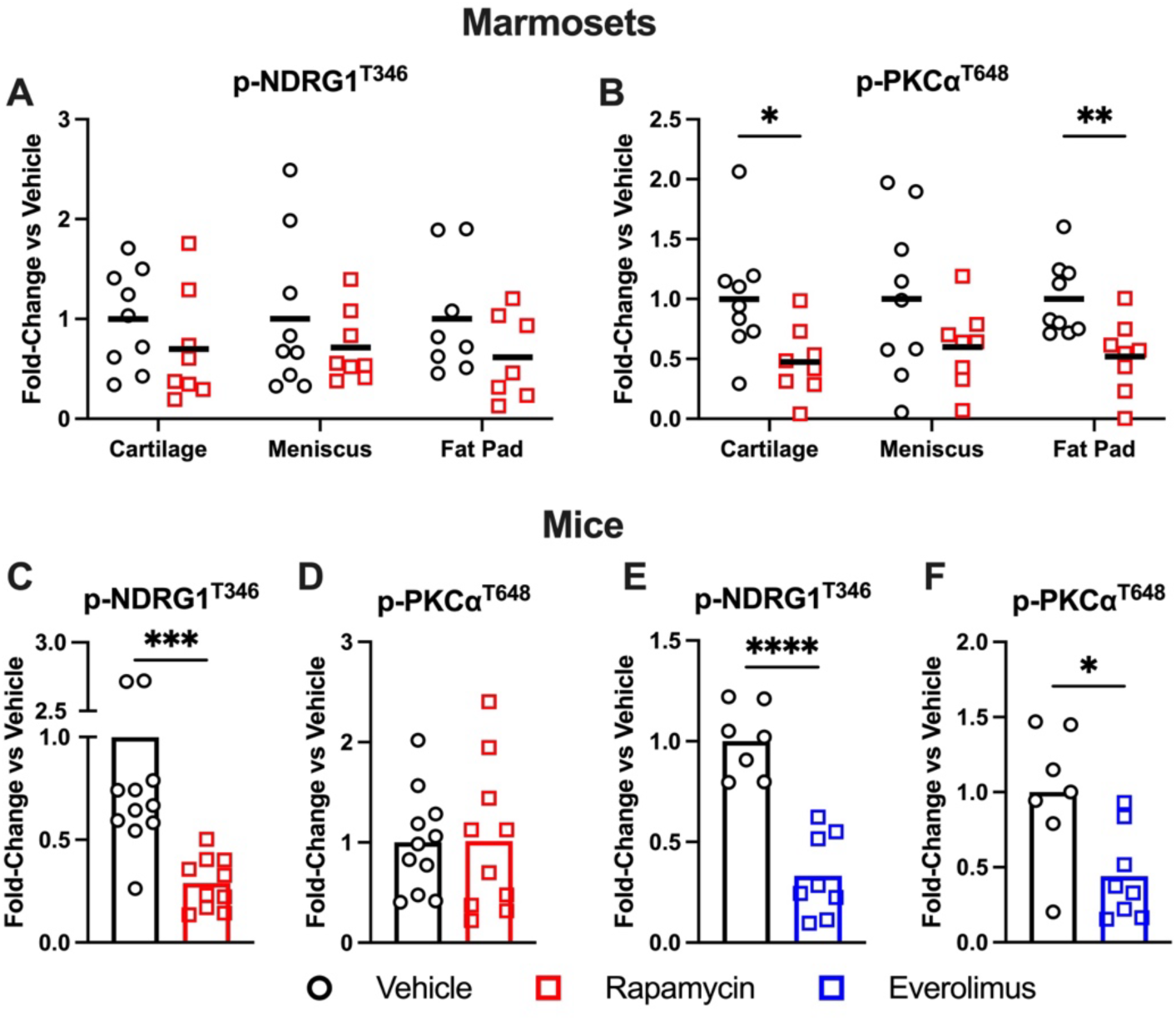
Alternative mTORC2 substrates are inhibited or unaffected during rapamycin treatment. **A**) Densitometric analysis of p-NDRG1^T346^ and **B**) p-PKCα^T638^ normalized to Ponceau S from control and rapamycin-treated marmoset articular cartilage, meniscus, and infra-patellar fat pad. **C**) Densitometric analysis of p-NDRG1^T346^ and **D**) p-PKCα^T638^ normalized to Ponceau S from glenohumeral cartilage of female C57BL/6J mice treated with rapamycin (2mg/kg, 3x/week) or vehicle for 8-weeks. **E**) Densitometric analysis of p-NDRG1^T346^ and **F**) p-PKCα^T638^ normalized to Ponceau S from glenohumeral cartilage of male mice treated with everolimus (3mg/kg every other day) or vehicle for 5-weeks. N=7-9/group (A-B), 10-12/group (C-D), or 7-8/group (E-F). Data are shown as mean with individual data points and were analyzed by student’s t-test. *P<0.05, **P<0.01, ***P<0.001, ****P<0.0001.

In glenohumeral cartilage from vehicle- and rapamycin-treated female mice, p-NDRG1^T346^ was significantly inhibited (**Figure 6C**) and p-PKCα^T638^ was unaffected by rapamycin (**Figure 6D**). In everolimus-treated male mice, both p-NDRG1^T346^ and p-PKCα^T638^ were significantly inhibited versus vehicle (**Figure 6E-F**). These data demonstrate that, unlike p-Akt^S473^, other mTORC2 substrates are not upregulated by rapamycin and rapamycin analogs in joint tissues.

### Low concentrations of rapamycin induce Akt^S473^ phosphorylation

There is evidence that small molecule delivery can be hindered by poor penetration and/or short residence time within joint tissues resulting in low effective concentration (29). Therefore, to investigate if these are factors which may be impacting molecular target engagement of rapamycin on mTOR signaling, we performed dose-response and time-course experiments using the chondrogenic ATDC5 cell line.

First, we treated ATDC5 cells with rapamycin doses ranging from 0.5nM to 100nM for 24-hours (**Figure 7A**). At all doses, p-RPS6^S235/236^ was completely diminished (**Figure 7B**). p-Akt^S473^ was increased ∼2-fold versus vehicle by 0.5nM rapamycin and returned to baseline levels at higher doses (**Figure 7C**). Next, we treated cells with 100nM rapamycin for 1- or 24-hours (**Figure 7D**). At both timepoints p-RPS6^S235/236^ was ablated by rapamycin (**Figure 7E**) and p-Akt^S473^ was unaffected (**Figure 7F**).

**Figure 7:**
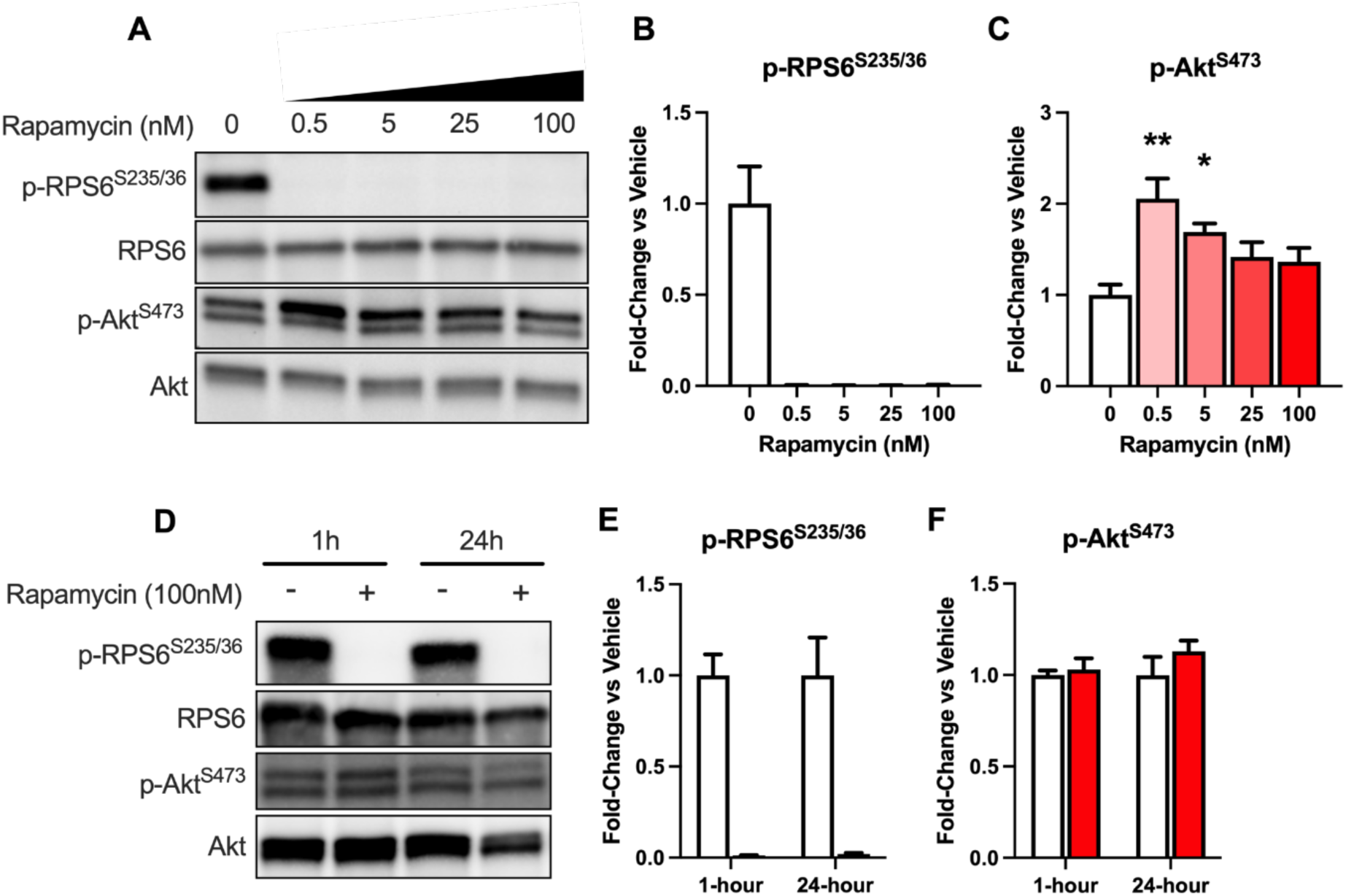
Low concentrations of rapamycin increased p-Akt^S473^ in ATDC5 cells. **A**) Representative western blots from ATDC5 cells treated with increasing doses of rapamycin for 24-hours. **B**) Densitometric quantification of p-RPS6^S235/236^ /total and **C**) p-Akt^S473^ /total. Data were analyzed via one-way ANOVA with Holm-Sidak’s multiple comparison test (B, C). *P<0.05, **P<0.01 vs. 0 nM rapamycin. **D**) Representative western blots from ATDC5 cells treated with 100nM rapamycin or DMSO vehicle for 1- or 24-hours. **E**) Densitometric quantification of p-RPS6^S235/236^ /total and **F**) p-Akt^S473^ /total. Data were analyzed using multiple t-tests (E-F). All data are presented as mean±SEM. N=3/group.

Conversely, low concentrations of rapamycin did not impact NDRG1 and PKCα phosphorylation, which were inhibited by chronic exposure to higher concentrations. Representative dose-response blots are shown in **Figure 8A**. At doses 5nM and higher, p-NDRG1^T346^ and p-PKCα^T638^ were decreased ∼40-50% versus vehicle, which only reached significance for NDRG1 (**Figure 8B-C**). Representative time course blots are shown in **Figure 8D** demonstrating p-NDRG1^T346^ was unaffected by 100nM rapamycin at 1-hour but was reduced ∼50% after 24-hours (**Figure 8E**). Together, these data indicate low concentrations of rapamycin produce similar molecular effects on mTORC1/2 signaling as were seen in mouse and common marmoset joint tissues.

**Figure 8:**
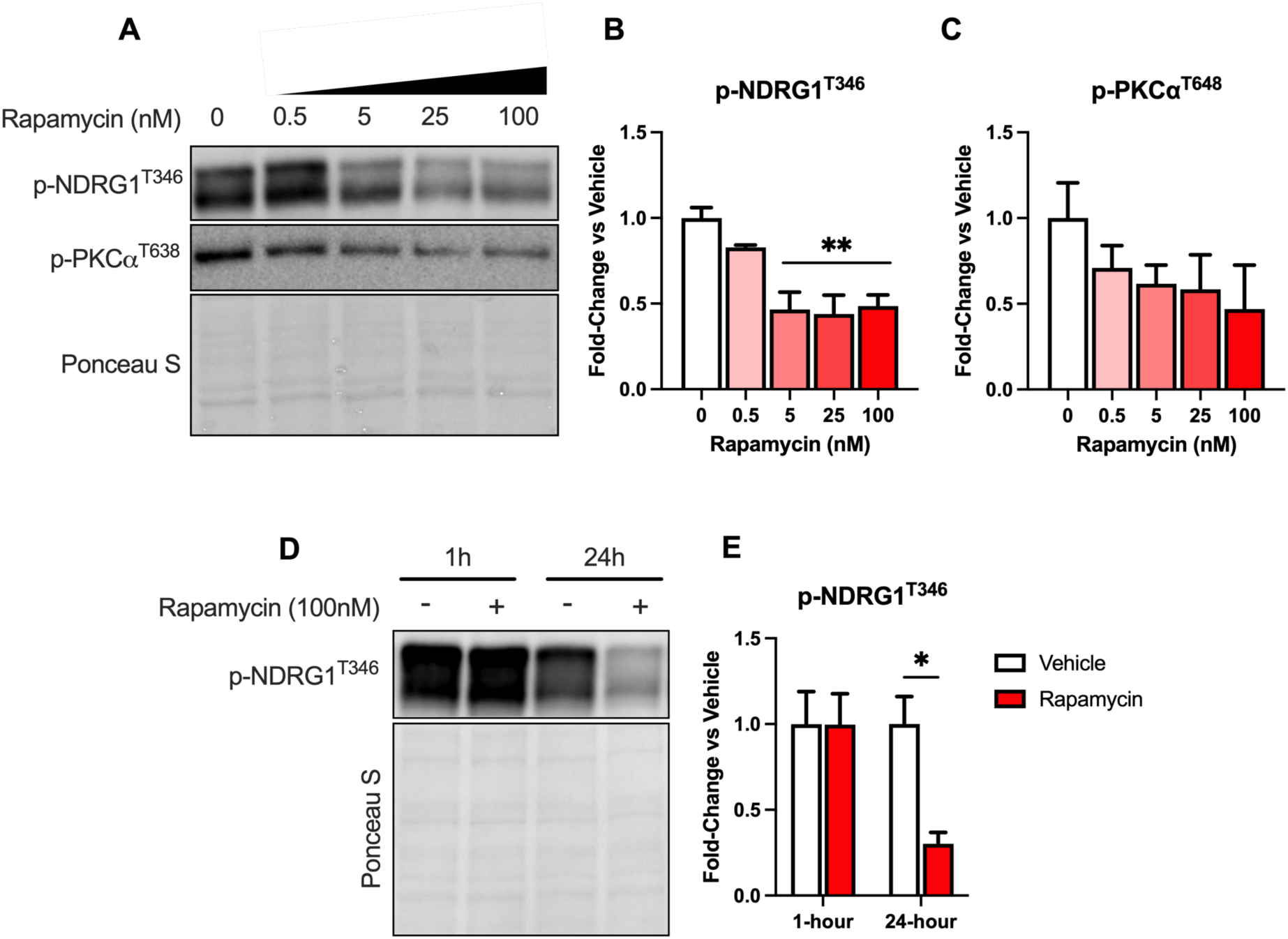
High doses of rapamycin and prolonged exposure decrease mTORC2 substrate phosphorylation. **A**) Representative western blots from ATDC5 cells treated with increasing doses of rapamycin for 24-hours. **B**) Densitometric quantification of p-NDRG1^T346^ /Ponceau S and **C)** p-T638-PKCα/total. Data were analyzed using one-way ANOVA with Holm-Sidak’s multiple comparison test (B-C). **P<0.01 vs. 0 nM rapamycin. **D**) Representative western blots from ATDC5 cells treated with 100nM rapamycin or DMSO vehicle for 1- or 24-hours. **E**) Densitometric quantification of p-NDRG1^T346^ /Ponceau S. *P<0.05 vs. vehicle. Data were analyzed using multiple t-tests. All data are presented as mean±SEM. N=3/group.

## DISCUSSION

Systemic rapamycin treatment worsened hallmarks of OA in female mice and marmosets. Specifically, rapamycin increased age-related meniscus calcification in female marmosets and exacerbated meniscus calcification and cartilage pathology after ACLR in female mice. In multiple joint tissues from mice and marmosets, rapamycin inhibited mTORC1 signaling as evident by lower p-RPS6^S235/236^ with feedback activation of p-Akt^S473^. Akt^S473^ is largely regulated by mTORC2 (30), however, other mTORC2 substrates were inhibited by mTOR-inhibitors, suggesting rapamycin and rapamycin analogs may differentially regulate downstream targets of mTORC2 or modify alternative upstream kinase activity that could contribute to Akt activation.

Upregulation of mTOR signaling during OA is conserved across multiple species including humans (7), and constitutively active mTORC1 activity can induce and exacerbate OA in mice (6). Genetic deletion of the entire mTOR kinase had the strongest protective effects against surgically induced OA (7), however, the impact of pharmacological mTOR-inhibition by rapamycin on OA has been more equivocal (6,7,11,12,31). We previously found 12-weeks of dietary rapamycin treatment (14ppm) with or without metformin co-treatment (1000ppm) worsened cartilage pathology in male Dunkin-Hartley guinea pigs (12). In models of temporomandibular joint (TMJ) OA, rapamycin (8mg/kg, IP, alternating days) induces chondrocyte primary cilia loss and an aged TMJ phenotype in young male mice (32), and exacerbates subchondral bone loss and TMJ OA in male rats (5mg/kg/day, intra-articular) (11).

Meniscus calcification was greater in female geriatric marmosets and ACLR-induced mice treated with rapamycin. This effect was not observed in male marmosets, though this may be due to their low propensity to develop meniscus calcification with age compared to females (20). In agreement, we found no effect of rapamycin on meniscus calcification in the contralateral limb of our ACLR-induced mice, suggesting this effect of rapamycin is from interaction with OA-associated processes. The role of mTOR in meniscus calcification is not well studied, however, rapamycin can enhance calcification during inflammatory conditions in bone marrow stem cells (BMSCs) *in vitro* and increases rat alveolar bone formation during inflammatory induction with lipopolysaccharides (33). These data would support the notion that within a pro-inflammatory environment, as commonly seen during OA, rapamycin treatment may create a condition prone to osteogenesis and aberrant calcification.

Subchondral bone thickness was reduced in the lateral tibia of geriatric rapamycin-treated marmosets. Interestingly, subchondral bone loss was not observed in other joint compartments. We previously found marmoset subchondral cortical and trabecular bone thickness increased with age in the medial but not lateral tibia and femur (20). Therefore, the reduction in lateral tibia subchondral thickness by rapamycin was not combatting an age-related increase and may be a maladaptive effect. We and others have previously observed decreased bone mass after rapamycin treatment. Twelve-weeks of dietary rapamycin in Dunkin-Hartley guinea pigs decreased cortical thickness in the medial joint compartment but also at other non-OA-affected sites (12). Further, rapamycin (4mg/kg every other day) decreased cortical and trabecular bone thickness and increased serum markers of bone resorption in 12-16-week-old female mice (34). However, the loss of bone mass during rapamycin treatment may be dependent on frequency of administration, as intermittent rapamycin (2mg/kg, every 5-days) does not impair bone mass maintenance in adult female mice (35).

Finally, we found the effects of rapamycin on mTOR signaling in marmoset and mouse joint tissues were conserved across species. In marmoset cartilage, meniscus, and infra-patellar fat pad, and in mouse glenohumeral cartilage, rapamycin treatment decreased relative phosphorylation of the mTORC1 effector RPS6^S235/236^ and increased phosphorylation of the mTORC2 effector Akt^S473^. We have previously shown in male Dunkin-Hartley guinea pigs, dietary rapamycin (14ppm) with or without metformin (±1000ppm) decreased mTORC1 but increased Akt^S473^ signaling when all rapamycin-treated animals were retrospectively pooled (due to close phenotypic similarity), and this was accompanied by worsened OA pathology (12). There is evidence that PI3K-Akt signaling promotes OA progression (28). Excessive activation of PI3K via PTEN-KO increases p-Akt^S473^ and can induce OA, and concurrent deletion of Akt alleviates these effects (5). Therefore, Akt-activation may be an important contributor to the deleterious effects of rapamycin observed in our present and previous studies (12). Because previous studies using rapamycin or other mTOR-inhibition strategies in OA models (6–8,36,37) have only characterized molecular target engagement with mTORC1 substrates, it is unclear whether activation of Akt and effects on other mTORC2 substrates are findings unique to our studies.

Rapamycin can have varying effects on mTORC2 signaling depending on cell and tissue type (38,39), including increased Akt phosphorylation. The mTORC1 effector S6K can inhibit upstream receptor tyrosine kinases as part of a negative feedback loop. During rapamycin treatment, mTORC1 is potently inhibited, alleviating this negative feedback loop and allowing activation of PI3K-Akt signaling (40). Akt^S473^ phosphorylation is canonically regulated by mTORC2 (30,38), however, we did not observe a similar increase in phosphorylation of other mTORC2 substrates such as p-NDRG1^T346^ and p-PKCα^T638^, which were unaffected or inhibited during rapamycin treatment. There is evidence of context-dependent substrate preference of mTORC2, depending on upstream signals and subcellular localization of mTORC2 (41,42). This may explain our varying observations across mTORC2 effectors. However, rapamycin or Torin-1 can induce Akt^S473^ phosphorylation in *Sin1^-/-^* or *Rictor^-/-^* mouse embryonic fibroblasts (MEFs) lacking functional mTORC2 (43,44), suggesting an mTORC2-independent mechanism of Akt phosphorylation during mTORC1 inhibition. This may be partially regulated through Akt auto-phosphorylation during sufficient pressure from PI3K (44), though other kinases such as IkB kinase epsilon (IKKe) and TANK binding kinase 1 (TBK1) that are capable of phosphorylating Akt^S473^ in the absence of mTORC2 in MEFs (43,45). Interestingly, IKKe and TBK1 are increased in OA joint tissues and can promote disease progression in mouse models of OA (46,47). Further, in cardiac mTOR-KO mice, increased TBK1 phosphorylation is associated with increased p-Akt^S473^ (45). Together this both implicates mTORC2 and opens the possibility of other upstream kinases as regulators of feedback activation of Akt during rapamycin treatment.

Therapeutic efficacy of small molecules for OA is often hindered by poor diffusion into joint tissues and/or low retention within the synovial joint (29). In agreement with this concept, we found *in vitro* that low concentrations of rapamycin closely replicated the effects of rapamycin on Akt^S473^ phosphorylation in joint tissues from rapamycin-treated mice, guinea pigs (12), and marmosets.

Therefore, some of the deleterious effects we observed may be due to a low effective dose received by joint tissues leading to elevated Akt^S473^ phosphorylation. Previous studies have shown that conjugation or encapsulation strategies that improve cartilage penetration and retention of intra-articularly administered rapamycin are associated with greater therapeutic efficacy than free rapamycin (48–51) though it is unclear whether this is through more potent or prolonged inhibition of mTORC1, or action on mTORC2/Akt signaling. In contrast to our findings, these studies report, at worst, no effect of unconjugated rapamycin on OA pathologies. Therefore, the adverse effects we observed may be due to the interaction of rapamycin with disease processes specific to the OA models employed. It will be important for future work to consider if the timing of mTOR-inhibition relative to the onset of OA as well as the rate of disease progression, dictate whether mTOR-inhibition is therapeutic or deleterious.

### Limitations

This work highlights important considerations for the efficacy of rapamycin as an OA therapeutic, however, we acknowledge several limitations. First, the primate study design was cross-sectional and included marmosets of varying ages. Because marmoset OA is highly related to age, there was likely pre-existing age-related decline in joint tissue integrity and/or OA pathology in some marmosets by middle to older age when treatment started, which is not reflective of a prophylactic treatment regimen. Being a genetically diverse primate, marmosets develop age-related OA in a variable fashion. While this may have limited our ability to detect more subtle effects of rapamycin on radiographic and histological OA, this heterogeneity is reflective of the human population. Further, despite this variability, we still observed robust treatment effects on subchondral bone and meniscus calcification in female marmosets which were corroborated in more controlled mouse experiments.

Our mTOR signaling outcomes for mouse experiments were performed on cartilage from the glenohumeral joint, which was not the site of OA induction. This was primarily due to antibody reactivity; we and, to our knowledge, no other investigators have been able to successfully detect p-Akt^S473^ via immunohistochemistry in mouse joint tissues, and major studies assessing Akt signaling in cartilage have also relied on western blot as the primary outcome (5). However, our mTOR signaling outcomes in marmosets were performed in knee joint tissues and closely mirrored our findings in mice. Finally, cell culture experiments investigating dose- and time-dependent effects of rapamycin were performed in a chondrogenic cell line. While they are not primary chondrocytes, ATDC5s have been previously used to investigate the impact of rapamycin and mTOR signaling on OA-associated mechanisms (6) and reflected our *in vivo* findings.

### Conclusions

In conclusion, we report several deleterious effects of rapamycin on OA-associated pathologies in female common marmosets and mice. These pathologies are associated with increased activation of Akt^S473^, which is consistent with exposure to low concentrations of rapamycin. Future work should investigate whether alternative dosing or delivery methods can prevent these deleterious effects of rapamycin by inhibiting both mTORC1 and Akt^S473^ and explore the role of other upstream kinases in rapamycin-induced Akt activation.

## ACKNOWLEDGEMENTS

This work is supported by supported by NIH-NIA R21-AG067464 (to ARK), American Physiological Society John F. Perkins Jr. Research Career Enhancement Award (to ARK), NIH-NIA R01-AG050797 (to ABS), the San Antonio Nathan Shock Center (P30-AG013319), and the San Antonio Claude D. Pepper Center (P30-AG044271). DMM is supported by NIH-NIA T32-AG000213. The Konopka Laboratory is also supported by NIH-NIA U01-AG076941, U01-AG081482, Impetus Longevity Grant by the Norn Group and Hevolution Foundation, and New Investigator Award from AFAR and Hevolution Foundation. The Lamming lab is supported in part by the NIA (AG056771, AG081482, AG084156 and AG085898), the NIDDK (DK125859), the Wisconsin Partnership Program, and startup funds from UW-Madison. MicroCT imaging was conducted at the Biomedical Imaging Center of the Beckman Institute for Advanced Science and Technology at the University of Illinois Urbana-Champaign and at the Small Animal Imaging and Radiotherapy Facility at the Wisconsin Institutes for Medical Research. Histology services were used through the University of Wisconsin Translational Research in Pathology Laboratory (TRIP) which is supported by the UW Department of Pathology and Laboratory Medicine, UWCCC (P30 CA014520), and the Office of The Director NIH (S10 OD023526). The Konopka Laboratory is also supported by startup and other funds from the University of Wisconsin-Madison School of Medicine and Public Health and Department of Medicine, and this work was supported using facilities and resources from the William S. Middleton Memorial Veterans Hospital. The authors would also like to acknowledge the lives of the animals used in the present study. The content is solely the responsibility of the authors and does not necessarily represent the official views of the NIH, the Department of Veterans Affairs, or the United States Government.

## Conflicts of Interests

DWL has received funding from, and is a scientific advisory board member of, Aeovian Pharmaceuticals, which seeks to develop novel, selective mTOR inhibitors for the treatment of various diseases.

## Notes

### Summary of Updates

Added additional mouse and in vitro experiments. Data are in new Figures 4 - 8.

