## Supplementary Information for "Inhibition of mTORC1 by rapamycin results in feedback activation of Akt^S473^ and aggravates hallmarks of osteoarthritis in female mice and non-human primates"

**METHODS**

All radiographic and histological analyses of marmoset joint tissues were performed as outlined in our previous manuscript (1), and all control and rapamycin-treated marmosets were processed and analyzed in parallel. Detailed methods were published in our previous manuscript but are replicated below for ease of readership:

microCT Acquisition and Quantitative Analysis

Knee joints were scanned while submerged in 70% ethanol using a Rigaku CT Lab GX130 at 120uA and 110kV for 14-minutes, achieving an isometric voxel size of 49um as previously performed (2). Three hydroxyapatite phantoms were included on each day of scanning to calculate bone mineral density.

For quantitative analysis, all scans were binarized using identical thresholds to allow for comparison across scans. In Amira (v2021.2, ThermoFisher), regions of calcified meniscus and were segmented, and volumes were quantified using the material statistics function. Representative 3D reconstructions were also created in Amira.

For analysis of subchondral bone, cortical and trabecular bone were separated using a previously validated algorithm (3). Three-dimensional thickness and density maps were created the ImageJ plug-in BoneJ. Articular surfaces were spatially normalized to account for morphological differences between marmosets, resulting in 81 subregions in the tibial plateaus and 40 subregions in the femoral condyles. For each sub-region, mean trabecular thickness and density were calculated. Subregions were then compiled to produce mean and max values for each joint compartment, as well as standard deviation as an index of heterogeneity of thickness and density across the articular surface. Three scans from the geriatric group could not be analyzed due to severe subchondral bone degeneration.

Semi-Quantitative microCT Scoring

To assess radiographic severity of OA, uCT scans were graded by a veterinary radiologist according to the criteria outlined in **Table S4** (4). Grading was performed in a single session utilizing standard transverse plane images and multiplanar reconstructions in sagittal and coronal planes while remaining blinded to marmoset age or sex. We also employed a semi-quantitative scoring system to evaluate calcification of the menisci. Briefly, the medial and lateral menisci were each assigned a score ranging from 0-5 depending on the size and frequency of ectopic calcification.

Histological Analysis

Following microCT scanning, knee joints were grossed of soft tissue, trimmed, and decalcified in 5% EDTA changed 2-3rx/week for 15-weeks or until fully decalcified by mechanical endpoint testing. Joints were then transected along the medial collateral ligament and processed for paraffin embedding. Sections were taken at 5-microns and stained with Toluidine Blue as previously described (1,2). In a blinded fashion, histological assessment of OA was performed using a grading scheme shown in **Table S5** which was adapted from the Modified Mankin guidelines for mice and other small rodents (5). Briefly, 3 sections across each joint compartment were graded in a blinded fashion for articular cartilage structural damage based on lesion depth and affected proportion of cartilage surface (0-11), tidemark duplication (0-3), proteoglycan loss as evident by loss of toluidine blue staining (0-8), and disruption of normal chondrocyte cellularity (0-4). Data were then presented as an average of the 3 scores. These criteria were assessed as individual phenomena and summed to produce a total histological score for each joint compartment and a whole-joint score.

### SUPPLEMENTARY TABLES

**Table S1: Modified OARSI scoring criteria.**

| Score | Observation |
|---|---|
| 0 | Intact, strong staining along the femoral condyle and tibia |
| 1 | Minor fibrillation without cartilage loss |
| 2 | Clefts below the superficial zone |
| 3 | Cartilage thinning on the femoral condyle and tibia |
| 4 | Lack of staining on the femoral condyle and tibia |
| 5 | Staining present on 90% of the entire femoral condyle with tibial degeneration |
| 6 | Staining present on 80% of the entire femoral condyle with tibial degeneration |
| 7 | Staining present on 75% of the entire femoral condyle with tibial degeneration |
| 8 | Staining present on 50% of the entire femoral condyle with tibial degeneration |
| 9 | Staining present on 25% of the entire femoral condyle with tibial degeneration |
| 10 | Staining present on <10% of the entire femoral condyle with tibial degeneration |

| | Male | | Female | | Pooled Sexes | |
|---|---|---|---|---|---|---|
| | N | Duration (yrs) | N | Duration (yrs) | N | Duration (yrs) |
| Adult | 3 | 0.6 (-0.15,1.35) | 2 | 1.3 (-1.39,4.0) | 5 | 0.9 (0.28,1.52) |
| Geriatric | 11 | 2.7 (1.63,3.78) | 8 | 2.2 (0.86,3.54) | 19 | 2.5 (1.72,3.27) |
| Pooled Ages | 14 | 2.2 (1.22,3.18) | 10 | 2.0 (1.0,3.0) | 24 | 2.1 (1.47,2.73) |

**Table S2**. Duration of rapamycin treatment in years for marmosets presented as pooled and stratified by age-group and sex. Number of marmosets is shown for each sub-group. Data are presented as mean (95% CI).

| Variable | Control | Rapamycin | P-value |
| --- | --- | --- | --- |
| Peak bodyweight (g) | 446.6 (424.4, 468.8) | 430.1 (401.2, 459.0) | 0.36 |
| Blood glucose (mg/dL) | 161.3 (136.3, 186.2) | 199 (136.0, 262.0) | 0.15 |

**Table S3**: Peak bodyweight, lean mass, fat mass, blood glucose, LDL cholesterol, and triglycerides are shown for control and rapamycin-treated marmosets. Data are presented at mean ± SD. Treatment groups were compared using 2-tailed, unpaired t-tests and P-values are presented.

**Table S4**: microCT scoring system for radiographic evaluation of OA.

| **MicroCT Finding** | **Score** |
|---|---|
| Osteophyte location | 1 = medial/lateral tibia<br>2 = patella<br>3 = medial/lateral femur |
| Osteophyte size | 0 = none<br>1 = small (<1mm)<br>3 = large (≥1mm) |
| Subchondral bone cystic changes | 0 = no<br>1 = yes |
| Subchondral bone sclerosis | 0 = no<br>1 = yes |
| Articular bone lysis | 0 = no<br>1 = yes |
| Intra-articular soft tissue | 0 = normal<br>1 = increased |

**Table S5**: Grading criteria for histological evaluation of OA in marmosets.

| Parameter | Grade | Description |
|---|---|---|
| Articular Cartilage Structure | 0 | Normal |
| | 1 | Undulating articular surface, but no fibrillation |
| | 2 | Mild superficial fibrillation involving < half of the plateau/condyle |
| | 3 | Mild superficial fibrillation involving > half of the plateau/condyle |
| | 4 | Mild fibrillation/clefts/loss involving up to 1/3 depth of articular cartilage thickness in < half of plateau/condyle |
| | 5 | Mild fibrillation/clefts/loss involving up to 1/3 depth of articular cartilage thickness in ≥ half of plateau/condyle |
| | 6 | Moderate fibrillation/clefts/loss involving up to 2/3 depth of articular cartilage thickness in < half of the plateau/condyle |
| | 7 | Moderate fibrillation/clefts/loss involving up to 2/3 depth of noncalcified articular cartilage thickness in & half of the plateau/condyle |
| | 8 | Severe fibrillation/clefts/loss involving > 2/3 depth of articular cartilage thickness in < half of plateau/condyle |
| | 9 | Severe fibrillation/clefts/loss involving > 2/3 depth of articular cartilage thickness in > half of plateau/condyle |
| | 10 | Clefts/loss of articular cartilage through tidemark |
| | 11 | Clefts/loss of articular cartilage through to subchondral bone |
| Tidemark Duplication | 0 | None (only one tidemark) |
| | 1 | 2 tidemarks |
| | 2 | >2 tidemarks |
| | 3 | No visible tidemark remaining |
| Toluidine Blue Staining | 0 | Normal (no loss of staining in articular cartilage) |
| | 1 | Moderate loss of staining in up to 1/2 depth of articular cartilage thickness and involving < half of the plateau/condyle |
| | 2 | Moderate loss of staining in up to 1/2 depth of articular cartilage thickness involving > half of the plateau/condyle |
| | 3 | Moderate loss of staining in > 1/2 depth of articular cartilage thickness and involving < half of plateau/condyle |

| | | |
|---|---|---|
| | 4 | Moderate loss of staining in > 1/2 depth of articular cartilage thickness and involving 2 half of plateau/condyle |
| | 5 | Severe loss of staining up to 1/2 depth of articular cartilage thickness and involving < half of plateau/condyle |
| | 6 | Severe loss of staining up to 1/2 depth of articular cartilage thickness and involving ≥ half of plateau/condyle |
| | 7 | Severe loss of staining in >1/2 depth of articular cartilage thickness and involving < half of the plateau/condyle |
| | 8 | Severe loss of staining in > 1/2 depth of articular cartilage thickness and involving ≥ half of plateau/condyle |
| Chondrocyte Cellularity | 0 | 0-5 groupings of hypertrophic chondrocytes or chondrones containing <3 cells. |
| | 1 | 6-10 groupings in chondrones or clusters containing 3 or more cells. |
| | 2 | >10 groupings of chondrones or clusters containing 3 or more cells. |
| | 3 | Loss of cellularity in <1/2 of the plateau or condyle. |
| | 4 | Loss of cellularity in 1/2 or greater of the plateau or condyle. |
